# Neutralization of viruses with European, South African, and United States SARS-CoV-2 variant spike proteins by convalescent sera and BNT162b2 mRNA vaccine-elicited antibodies

**DOI:** 10.1101/2021.02.05.430003

**Authors:** Takuya Tada, Belinda M. Dcosta, Marie Samanovic-Golden, Ramin S. Herati, Amber Cornelius, Mark J. Mulligan, Nathaniel R. Landau

## Abstract

The increasing prevalence of SARS-CoV-2 variants with mutations in the spike protein has raised concerns that recovered individuals may not be protected from reinfection and that current vaccines will become less effective. The B.1.1.7 isolate identified in the United Kingdom and B.1.351 isolate identified in the Republic of South Africa encode spike proteins with multiple mutations in the S1 and S2 subunits. In addition, variants have been identified in Columbus, Ohio (COH.20G/677H), Europe (20A.EU2) and in domesticated minks. Analysis by antibody neutralization of pseudotyped viruses showed that convalescent sera from patients infected prior to the emergence of the variant viruses neutralized viruses with the B.1.1.7, B.1.351, COH.20G/677H Columbus Ohio, 20A.EU2 Europe and mink cluster 5 spike proteins with only a minor decrease in titer compared to that of the earlier D614G spike protein. Serum specimens from individuals vaccinated with the BNT162b2 mRNA vaccine neutralized D614G virus with titers that were on average 7-fold greater than convalescent sera. Vaccine elicited antibodies neutralized virus with the B.1.1.7 spike protein with titers similar to D614G virus and neutralized virus with the B.1.351 spike with, on average, a 3-fold reduction in titer (1:500), a titer that was still higher than the average titer with which convalescent sera neutralized D614G (1:139). The reduction in titer was attributable to the E484K mutation in the RBD. The B.1.1.7 and B.1.351 viruses were not more infectious than D614G on ACE2.293T cells *in vitro* but N501Y, an ACE2 contacting residue present in the B.1.1.7, B.1.351 and COH.20G/677H spike proteins caused higher affinity binding to ACE2, likely contributing to their increased transmissibility. These findings suggest that antibodies elicited by primary infection and by the BNT162b2 mRNA vaccine are likely to maintain protective efficacy against B.1.1.7 and most other variants but that the partial resistance of virus with the B.1.351 spike protein could render some individuals less well protected, supporting a rationale for the development of modified vaccines containing E484K.

## Introduction

Since the zoonotic transfer of SARS-CoV-2 to humans at the end of 2019, the virus has rapidly mutated to adapt to its new host. Such adaptations are a feature of viral zoonosis in which selective pressure drives viral proteins to be optimized for interaction with the host cell proteins of the new species. In addition, viral amino acid sequences are selected to escape the humoral and cellular adaptive immune responses which recognize a different set of epitopes. While all viral genes are subjected to evolutionary pressure, the viral envelope glycoprotein is selected both for optimal interaction with its cell surface receptor and for escape from neutralizing antibodies.

In January, 2020, a variant SARS-CoV-2 was identified in Germany and China with a D614G mutation^1^ as compared to the presumptive zoonotic Wuhan isolate. By May, the variant had risen to a prevalence of >97% world-wide. Amino acid residue G614 lies in subdomain 2 near the S1:S2 processing site of the spike protein. The mutation was found to reduce S1 subunit shedding from virions leading to increased infectivity^2–4^ and the viral isolate replicated to higher titers in the upper respiratory tract although was not associated with increased mortality. Additional variants with increasing prevalence were identified. The B.1.1.7 lineage (VOC-202012/01) isolate which was identified in South East England, London and east of England^5–7^, was found to replicate with increased virus loads and to be increasing in prevalence with mathematical modeling suggested a 56% increase in transmissibility^6^. The B.1.1.7 lineage is defined by 23 mutations of which 8 are located in S (Δ69-70, Y144Del, N501Y, A570D, P681H, T716I, S982A and D1118H). N501Y is one of six contact residues with ACE2^8^ and has been shown to increase affinity for ACE2^9^. The Δ69-70, which was found in multiple independent lineages, increases viral infectivity and leads to immunoevasion in immunocompromised patients^10^. The P681H mutation lies adjacent to the furin cleavage site suggesting a possible role in spike protein processing.

In October, 2020, the B.1.351 lineage variant was identified in South Africa where it rapidly became the predominant circulating genotype^11^. The variant is more heavily mutated than B.1.1.7 with 9 mutations (L18F, D80A, D215G, L242-244del, R246I, K417N, E484K, N501Y and A701V) three of which (K417N, E484K and N501Y) are in the receptor binding domain (RBD). E484 and N501Y lie in the amino acid motif that directly contacts specific ACE2 residues. N501Y has been shown to enhance affinity of the spike protein for ACE2 by hydrogen bonding with ACE2 Y41 and is selected in a mouse model^12^. K417N, while not contributing to ACE2 binding, is a key epitope for the binding of neutralizing antibodies, as is E484K, and thus these mutations may have been selected for evasion of the humoral response^13–17^. Based on phylogenetic tree branch-length, it has been suggested that the variant arose through the prolonged virus replication in an immunocompromised individual^10^.

Additional variants found to be circulating in the human population include 20A.EU2 which was identified in Spain^18^ and later elsewhere in Europe. COH.20G/677H which was identified in Columbus, Ohio contains D614G, N501Y, Q677H but lacks the mutations present in the UK and South Africa variants suggesting an independent origin^19^. In addition, isolates with variant spike proteins were found in domesticated minks in Denmark with the potential for transfer into humans^20^.

The rapid emergence of new viral variants is of concern both because of their increased transmissibility and the possibility that they may escape neutralizing antibodies. Immunoevasion would have the potential to allow for the re-infection of individuals who have cleared the virus and could reduce the level of protection provided by currently approved vaccines and those in clinical trials which are based on the earlier viral isolate sequence. These include Pfizer BioNtech BNT162b2 lipid-nanoparticle-formulated, modified nucleoside mRNA vaccine encoding a trimerized spike protein^21^ and the Moderna mRNA-1273 vaccine that encodes the full-length stabilized spike protein^22^.

We report here on antibody neutralization of the European, UK, South Africa, Europe, Columbus, Ohio and mink spike variants by the sera of convalescent individuals and those vaccinated with BNT162b2. Neutralization was determined using lentiviral virions pseudotyped by the variant spike proteins. In addition, the infectivity, thermostability and affinity of the virions for ACE2 binding was determined. The results showed that the variants bound ACE2 with increased affinity and thermostability. The variants were neutralized both by the sera of convalescent and vaccinated individuals. Virus with the UK B.1.1.7 spike protein was neutralized as well as the parental D614G while the South Africa B.1.351 variant spike was neutralized with a 3-4-fold reduction in IC_50_ compared to D614G virus. Even with the relatively minor reduction in neutralizing titer for B.1.351 by serum antibodies of vaccinated individuals, neutralizing titers remained above those of naturally infected individuals. The findings suggest that the protection provided by vaccination will remain largely intact against the South Africa variant and other currently circulating SARS-CoV-2 variants. The partial resistance of B.1.351 to vaccine elicited antibody could result in reduced level of protection in individuals with below average neutralizing titer.

## Results

The trimeric SARS-CoV-2 spike protein is synthesized as a full-length precursor polypeptide that is processed by cellular proteases into S1 and S2 subunits (**Fig 1A**). S1, which mediates cell attachment, consists of an amino-terminal domain, the RBD, a receptor binding motif (RBM) within the RBD that directly contacts ACE2, and subdomains 1 and 2. S2, which mediates membrane fusion, consists of a hydrophobic amino-terminal fusion peptide, heptad repeats 1 and 2, a hydrophobic transmembrane domain and a cytoplasmic tail. The location of point mutations and small deletions in the major SARS-CoV-2 variant spike proteins are shown (**Fig 1A**). Mutations of concern are those lying in the RBD which is the binding site for most neutralizing antibodies and those within the RBM (453, 477, 484, 501) which directly contacts ACE2. Two mutations lie near the processing site and others are in the S1 NTD and S2. Whether the mutations act independently or in a coordinated fashion is not known and whether they were selected or are simply markers is not clear.

**Fig. 1.**
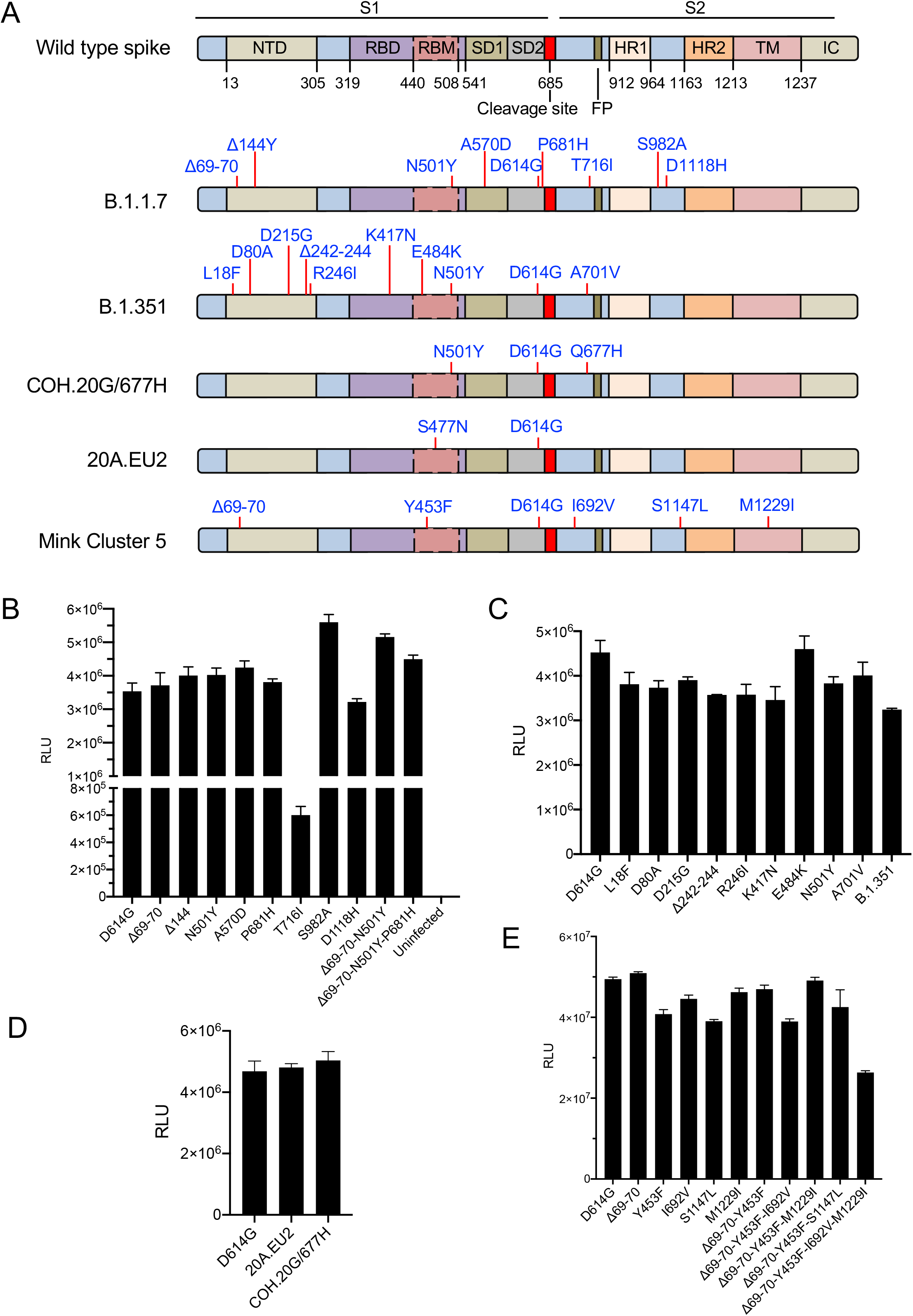
Infectivity of variant spike protein pseudotyped virus. (A) A schematic of the overall topology of the SARS-CoV-2 spike monomer is shown. NTD, N-terminal domain; RBD, receptor-binding domain; RBM, receptor-binding motif; SD1 subdomain 1; SD2, subdomain 2; FP, fusion peptide; HR1, heptad repeat 1; HR2, heptad repeat 2; TM, transmembrane region; IC, intracellular domain. The diagrams below show the location of mutations found in the UK variant B.1.1.7, South Africa variant B.1.351, Columbus, Ohio variant COH.20G/677H, European variant 20A.EU2, and mink cluster 5 variant. Spike protein expression vectors used to generate pseudotyped viruses were deleted for the carboxy-terminal 19 amino acids. (B) Lentiviral pseudotypes with individual mutations of the B.1.1.7 spike protein or combinations thereof were prepared. Infectivity of the virions normalized for RT activity was tested on ACE2.293T cells. Luciferase activity was measured two days post-infection. (C) Infectivity of virus with the South Africa B.1.351 variant spike protein and its component mutations. (D) Infectivity of virus with the Columbus, Ohio variant COH.20G/677H and European variant 20A.EU2 spike protein. (E) Infectivity of virus with the mink-associated variant spike protein individual mutations and combinations thereof. The experiments were repeated three times with similar results.

### Efficient pseudotyping of lentiviruses by variant spike proteins

Lentiviral pseudotypes provide a rapid and accurate means to assess spike protein function. Neutralizing antibody titers determined by lentiviral pseudotype assay closely mirror those measured by live SARS-CoV-2 assay^23^. To analyze the functional properties of the spike protein variants, we constructed cytomegalovirus (CMV) promoter-driven expression plasmids encoding the B.1.1.7, B.1.351, COH.20G/677H, 20A.EU2 and mink cluster 5 spike proteins or the component mutations, singly and in combination. Vector coding sequences were based on the Wuhan Hu-1 S gene with a deletion of the carboxy-terminal 19 amino acids that increases virion **(Fig. 1B)**. In this study, the D614G spike protein is considered “wild-type” and the variants tested contain G614. The expression vectors were used to generate lentiviral pseudotypes with a packaged genomic RNA encoding both GFP and nanoluciferase. Immunoblot analysis of the spike proteins showed that each was expressed in cells and incorporated into virions at levels comparable to wild-type D614G spike protein, with the exception of T716I and the fully mutated B.1.1.7 spike proteins which were expressed at lower levels (**Supplementary Fig. 1A and C**). Because the B.1.1.7 spike protein with the full complement of mutations was poorly expressed, we used the triple mutant Δ69-70/N501Y/P681H which contains the key B.1.1.7 mutations for neutralization experiments. Measurement of spike protein proteolytic processing as determined by the ratio of processed (S2) to full-length protein (S) showed that some of the mutations in the B.1.1.7 spike protein increased the extent of processing (N501Y, A570D, P681H, T716I, Δ69-70/N501Y and Δ69-70/N501Y/P681H) **(Supplementary Fig. 1A)**. B.1.351 mutations did not show an effect on processing **(Supplementary Fig. 1B)** as was also the case for the 20A.EU2 and COH.20G/677H spike proteins **(Supplementary Fig. 1D).**

The infectivity of virus with each of the variant spike proteins was tested by infection of ACE2.293T cells, a cell-line that expresses high levels of ACE2 with normalized amounts of pseudotyped viruses. Analysis of the B.1.1.7 variant and its component mutations showed that the single point mutations had little effect on infectivity (Δ69-70, Y144Del, N501Y, A570D, P681H and D1118H) except for T716I which was low and S982A which significantly increased infectivity **(Fig. 1B)**. The double mutant Δ69-70/N501Y and triple mutant Δ69-70/N501Y/P681H also had increased infectivity, suggesting that the two mutations coordinate to increase infectivity. Analysis of the B.1.351 proteins showed that the individual point mutations had similar infectivity while the full complement was slightly reduced **(Fig. 1C)**. Spike proteins with the mink-associated mutations were fully infectious except for the protein containing all 4 mutations (Δ69-70/Y453F/I692V/M1229F) which was 2-fold reduced in infectivity. The COH.20G/677H and 20A.EU2 variants were fully infectious **(Fig. 1D)**.

To determine whether differences in infectivity could be caused by effects of the mutations on the stability of the spike proteins or their incorporation into virions, we analyzed the spike proteins produced in transfected cell lysates and incorporated into virions. The transfected cell lysates were analyzed on an immunoblot probed for the spike protein S2, which allows for detection of the full-length spike protein and processed S1 protein, and for the HIV-1 capsid protein P24 as a means of normalizing for particle number. Analysis of B.1.1.7 showed that each of singly mutated proteins was expressed in cells at similar levels and processed to a similar extent **(Supplementary Fig. 1A)**. In contrast, some of the point mutations appeared to affect amount of spike on the virion and the extent of spike protein processing. T716I was expressed at significantly lower level, accounting for the decreased infectivity of this spike protein. P681H was present at high copy number but processed more efficiently than wild-type as demonstrated by a lower ratio of full-length to S2 protein level. The N501Y and A570D mutations resulted in a small decrease in copy number on virions. When combined with Δ69-70, processing returned to wild-type level. Addition of the P681H mutation in the triple mutant increased processing to a level similar to that of the P681H single point mutation. Analysis of the B.1.1.7 and B.1.351 spike proteins containing the full complement of mutations showed that the B.1.351 spike protein was stable and processed as wild-type while the B.1.1.7 was poorly expressed and little of the protein was present on virions **(Supplementary Fig. 1C)**.

### Convalescent sera neutralize UK and South Africa variants

A major factor in determining how well recovered patients are protected from re-infection with variant viruses is the extent to which antibodies elicited in primary infection cross-react on newly emerging viral variants. To determine the extent of antibody cross-reactivity, we tested neutralizing antibody titers of the serum specimens from 10 convalescent patients who had been infected prior to April, 2020, a period prior to the emergence of the viral variants, for neutralizing titers on the variant spike proteins. Viruses containing the point mutations of B.1.1.7 showed that the single point mutations (Δ69-70 and N501Y) were neutralized as efficiently as D614G. Spike protein with the combination of B.1.1.7 mutations (Δ69-70/N501Y/P681H) and the 20A.EU2 spike were neutralized slightly less well than D614G and this was noticeable in the lack of sera with high neutralizing titer for the viruses **(Fig. 2A)**. The Columbus, Ohio variant virus COH.20G/677H was neutralized with similar titer to D614G. A direct comparison of neutralizing titers of the B.1.1.7 spike mutations (Δ69-70, N501Y and Δ69-70/N501Y/P681H) with the D614G spike showed a close correlation of neutralizing titer for each donor **(Fig. 2B)**. This was also the case for COH.20G/677H, 20A.EU2 and mink cluster 5 spike proteins which were more easily neutralized than D614G virus. A detailed analysis of one donor serum chosen at random showed that the sera neutralized B.1.1.7 and its constitutive point mutations similarly, with the exception of T716I that was more easily neutralized than D614G **(Supplementary Fig. 2)**.

**Fig. 2.**
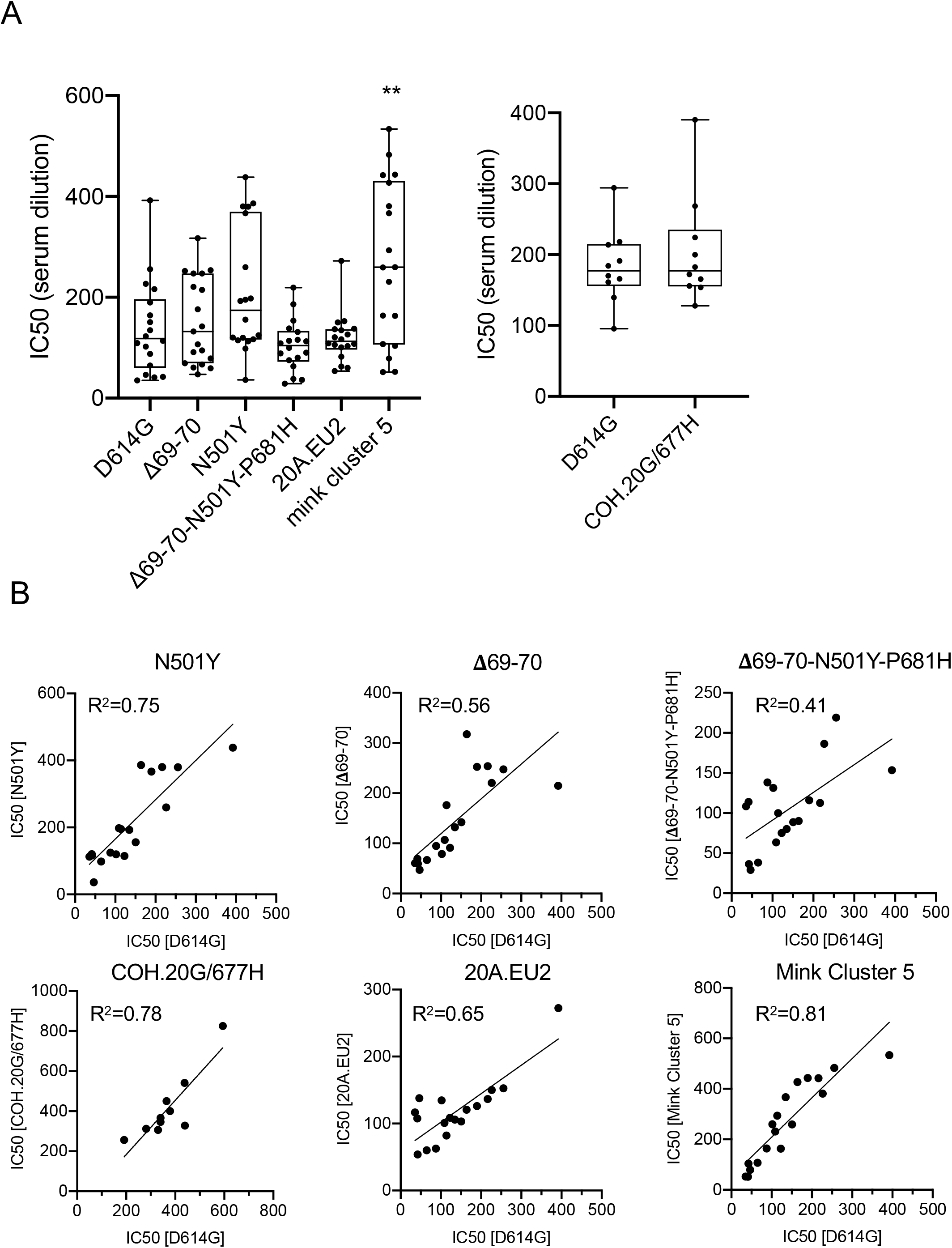
Neutralization of spike protein variant viruses by convalescent sera. (A) Serum samples from 18 (left) and 10 (right) convalescent individuals were serially diluted and incubated for 30 min with the lentiviral virions pseudotyped by the variant spike protein or by D614G ancestral spike protein and infectivity in ACE2.293T cells was then measured. The data represents serum neutralization IC50 of D614G and variant spike protein viruses. (B) The IC50s calculated from neutralization of N501Y, Δ69-70, Δ69-70-N501Y-P681H, COH.20G/677H, 20A.EU2, mink cluster 5 variants compared to D614G. Correlation analyses were performed using GraphPad Prism 8.

Analysis of B.1.351 and its constituent E484K point mutation showed that both viruses were neutralized by convalescent sera with titers similar to that of D614G **(Fig. 3A and B)**. For some donors, the neutralization curves were virtually identical (donors 1, 4, 5, 7, and 8) and for others the variant viruses were neutralized with slight reduction in titer (donors 2, 3, 6, 9 and 10). Overall the reduction in IC_50_ of B.1.351 and E484K viruses was about 1.7-fold (**Fig. 3C)**. IC_50_ for each donor for the E484K single mutant was similar to that of B.1.351 suggesting that the E484K single amino acid change was responsible for the decrease in neutralizing titer **(Fig. 3D)**. Spike proteins containing each of the other individual B.1.351 mutations were neutralized were neutralized as well as D614G (Supplementary Fig. 3).

**Fig. 3.**
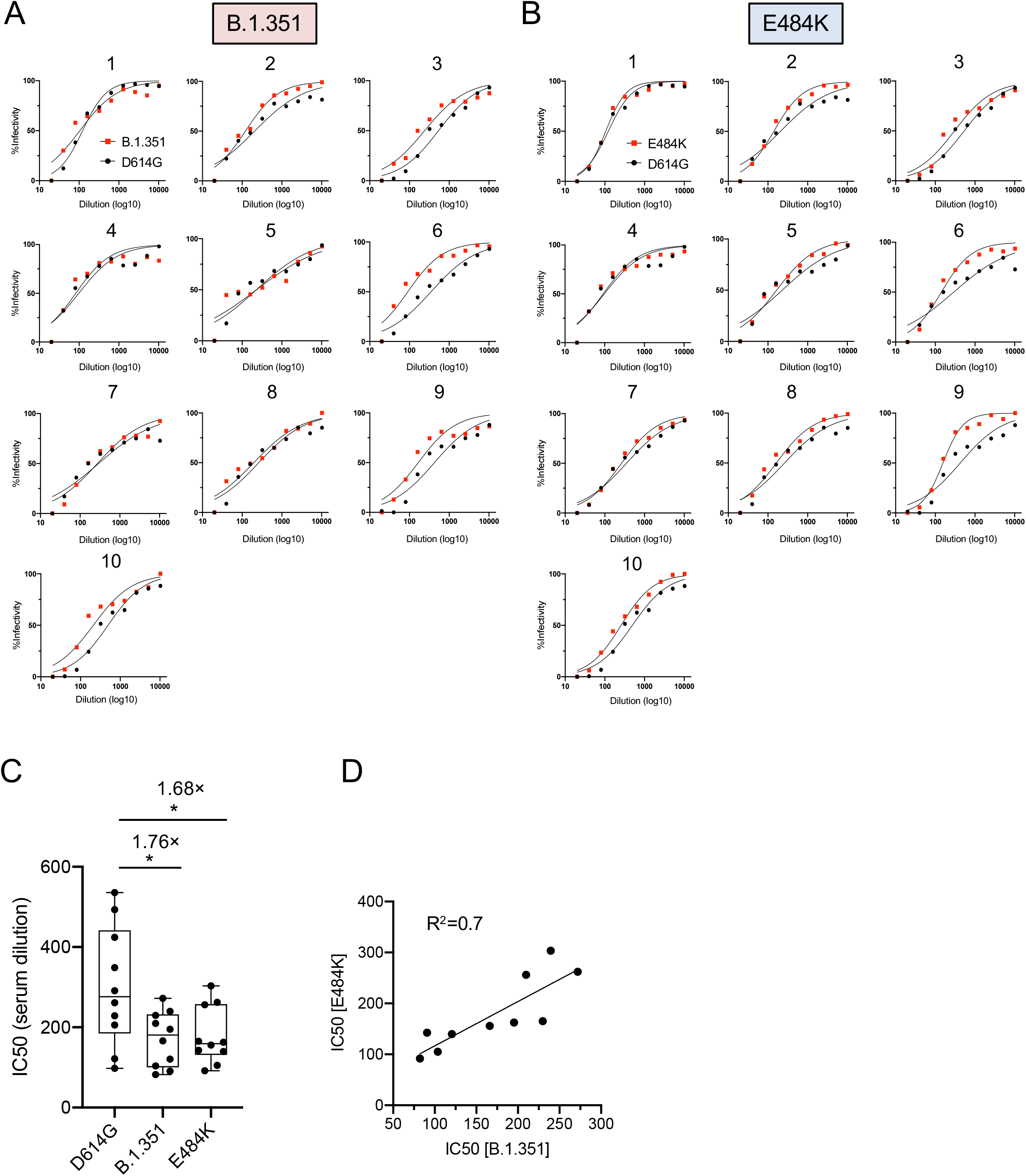
Viruses with B.1.351 and E484K pseudotyped virus are neutralized by convalescent sera. (A) Neutralization of virus with the South Africa B.1.351 pseudotyped virus by sera of 10 convalescent individuals. The data represents percent infectivity not neutralized by convalescent serum. (B) Serum neutralization curves of virus with the E484K spike protein mutation. (C) Neutralization IC50 of virus containing D614G, B.1.351 or E484K spike proteins by convalescent sera. (D) Comparison of neutralization IC50 of virus containing B.1.351 spike against E484K spike. Correlation analyses were performed using GraphPad Prism 8.

### Antibodies elicited by BNT162b2 vaccination neutralize viruses with the UK, South Africa, European and Columbus, Ohio spike proteins

The efficacy of current SARS-CoV-2 vaccines, which are based on spike proteins present prior to the emergence of viral variants, to protect against SARS-CoV-2 variants will be determined by how well the vaccine-elicited antibodies cross-react with circulating viral variants. To address this question, we analyzed the neutralizing activity of serum specimens from individuals vaccinated with the Pfizer mRNA vaccine against viruses with the B.1.1.7 and B.1.351 spike proteins. Sera were collected from five healthy donors on days 0, 7 and 28 where day 28 corresponded to 7 days post-second vaccine injection. At days 0 and 7, no detectable neutralizing antibody was detected indicating that the donors had not been previously infected (not shown). On day 28, all donors had high titers of neutralizing antibody against virus with the D614G spike protein with an average neutralizing titer of 1:1800, 7-fold higher than that of convalescent serum samples **(Fig. 4A and B)**. Neutralization titers against N501Y, S982A, B.1.1.7 (Δ69-70/N501Y/P681H), COH.20G/677H and 20A.EU2 were similar to D614G **(Fig. 4)** while B.1.351 and E484K were neutralized with a 3.1- and 4.3-fold decrease in titer. While this is a significant drop in titer, it remains higher than the titer found for convalescent sera against the D614G virus. The 3.1-fold decrease in neutralization of B.1.351 appears largely due to the E484K mutation **(Fig. 4C)**.

**Fig. 4.**
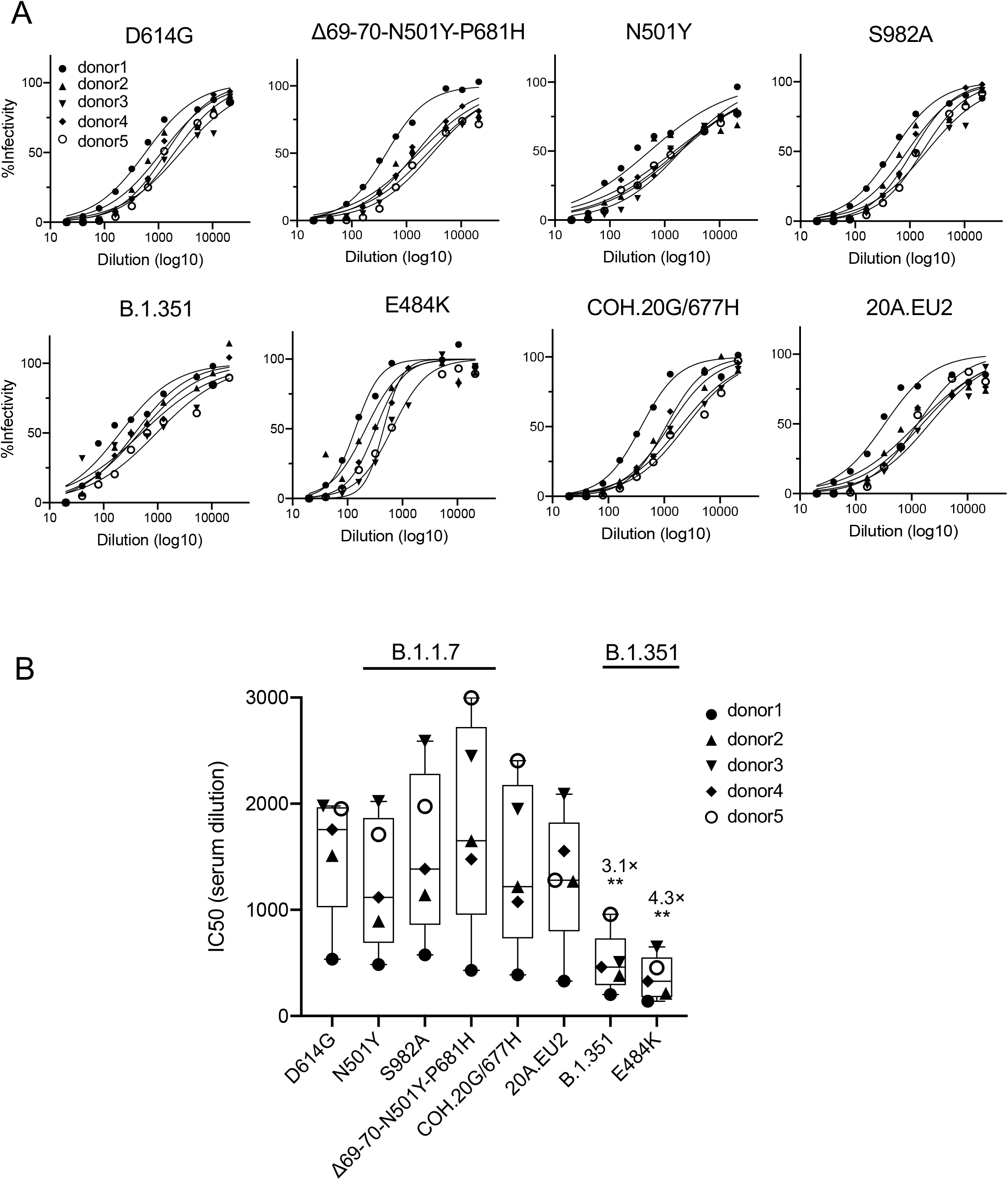
UK and South Africa variant spike proteins are neutralized by the sera of individuals who received BNT162b2 vaccine. (A) Serum samples from 5 vaccinated individuals were serially diluted and incubated for 30 min with lentiviral virions pseudotyped by the variant spike protein or by D614G ancestral spike protein and infectivity in ACE2.293T cells was then measured. The data represents percent infectivity not neutralized by serum. (B) Neutralization IC50 of viruses D614G, N501Y, S982A, Δ69-70/N501Y/P681H, COH.20G/677H, 20A.EU2 and B.1.351.

### Variant spike proteins have higher affinity for ACE2 and increased thermostability

The D614G mutation, which was first identified in January, 2020, caused a significant increase in viral infectivity and binding to ACE2. We previously reported on increased binding using an *in vitro* assay in which spike protein-pseudotyped virus was incubated with beads coated with soluble ACE2 (sACE2)^25^. Here, we used a more sensitive assay in which the pseudotyped viruses were incubated with free sACE2 and then allowed to infect ACE2.293T cells. In this assay, an increased sensitivity to sACE2 neutralization indicated higher affinity of the spike protein for ACE2. An analysis of viruses by this approach showed that the B.1.1.7 spike protein itself did not cause an increase in ACE2 binding; however, spike proteins containing the single N501Y mutation, as well as those that included N501Y (Δ69-70/N501Y and Δ69-70/N501Y/P681H), showed a significant increase in ACE2 binding compared to D614G **(Fig. 5A)**. Analysis of B.1.351 showed that it had increased ACE2 binding **(Fig. 5A)**. The increase was due to N501Y as none of the other point mutations had an effect. The COH.20G/677H spike protein, which also contains N501Y, displayed increased ACE2 binding **(Fig. 5C)**. Analysis of the proteins in the ACE2 binding assay in which virions were incubated with matrix-bound sACE2 confirmed that those spike proteins that contained N501Y had increased affinity for ACE2 (N501Y, Δ69-70/N501Y, COH.20G/677H and B.1.351). (**Fig. 5D**)

**Fig. 5.**
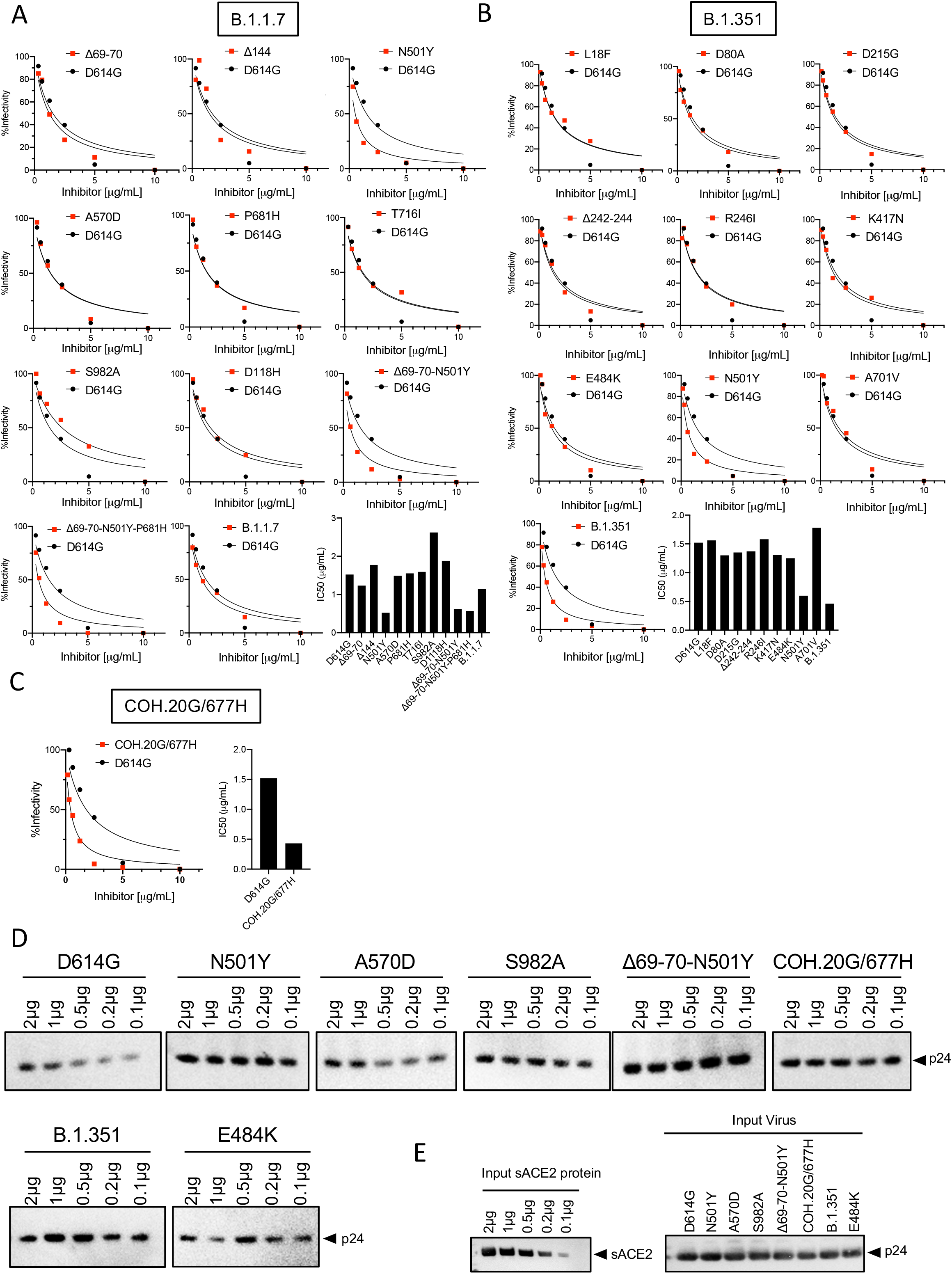
The effect of B.1.1.7 and B.1.351 spike protein mutations on ACE2 binding. The relative affinity of variant spike protein for ACE2 was measured in a soluble ACE2 (sACE2) neutralization assay (A-D) and virion binding assay (E). For sACE2 neutralization, virus containing variant protein was incubated for 30 min with serially diluted recombinant sACE2 and then used to infect ACE2.293T cells. After 2 days, luciferase activity was measured. The data represents percent infectivity, at varying concentrations of sACE2, of the spike variants and control D614G virus. For virion binding assay, pseudotyped virions (30 ng p24) were incubated for 30 min with Ni-NTA beads coated with the indicated amounts of His-tagged recombinant sACE2 protein. Unbound virions were removed by brief centrifugation and bound virions were analyzed on an immunoblot probed with anti-p24 antibody. (A) sACE2 neutralization of viruses with B.1.1.7 spike protein mutations. IC50s calculated from the neutralization curves are indicated. (B) sACE2 neutralization of viruses with B.1.351 spike protein mutations. IC50s calculated from the neutralization curves are indicated. (C) sACE2 neutralization of virus with the COH.20G/677H spike protein. IC50s calculated from the neutralization curves are indicated. (D) Virion binding assay of viruses with variant spike proteins. (E) Input sACE2 (left) and virus (right) for virus-sACE2 binding assay.

Transmissibility of the viruses is likely to be affected by their stability in aerosols and resistance to high temperature. A test of heat resistance over a short time period of viruses with spike proteins containing the critical mutations showed a gradual decrease in infectivity over 1 hour with increasing temperature **(Supplementary Fig. 4)**. Incubation of the virus for 1 hour at 50°C caused a 40-fold decrease in infectivity of the D614G virus. Viruses with N501Y, S982A, Δ69-70/N501Y/P681H spikes and B.1.351 decreased their infectivity less than 20-fold, suggesting that the mutations increase spike protein stability.

## Discussion

The emergence of SARS-CoV-2 variants with mutations in the spike protein has raised concerns as to whether antibodies elicited by primary infection and by vaccination will remain protective. We show that convalescent sera from individuals who had been infected prior to the emergence of the variants, neutralized viruses with the UK B.1.1.7, South Africa B.1.351, Columbus, Ohio COH.20G/677H, Europe 20A.EU2 and mink cluster 5 spike with titers similar to that of the earlier D614G spike protein. Sera of individuals vaccinated with the BNT162b2 mRNA vaccine neutralized D614G virus with an average titer 7-fold higher than convalescent sera. The vaccine-elicited antibodies neutralized virus with the B.1.1.7 spike protein with titers similar to D614G virus, suggesting high level protection from this variant. The vaccine-elicited antibodies neutralized virus with the B.1.351 spike with an average 3-fold reduction in titer (1:500), a titer that was still higher than that elicited by natural infection for virus with the D614G spike (1:139). The reduction in titer against the B.1.351 spike protein was attributable to the E484K mutation. Viruses with the B.1.1.7 and B.1.351 were not more infectious than those with D614G on ACE2.293T cells but N501Y, an ACE2 contact residue present in the B.1.1.7, B.1.351 and COH.20G/677H spike proteins, bound to ACE2 with increased affinity, likely contributing to the increased transmissibility of viruses with these spike proteins.

The variant spike proteins tested here were expressed well in cells and were efficiently incorporated into virions with the exception of the B.1.1.7 spike protein and one of its constituent mutations T716I. T716I, which is located close to the fusion peptide, caused a 7-fold decrease in infectivity and was incorporated at lower copy number into virions. The B.1.1.7 spike protein was present in cells at lower copy number and was incorporated into virions at lower copy number resulting in an 18-fold decrease in infectivity. Restoration of the amino acid 716 to threonine in the B.1.1.7 spike protein did not restore infectivity (not shown). This property may be specific to spike protein biosynthesis in 293T cells and is not likely to reflect the situation *in vivo* given the reported increase in infectivity of the virus. The pseudotyped virus retained ample infectivity to provide significant data in the neutralization assays.

Mutations in the spike protein could serve to increase infectivity of the virus, either by increasing affinity for ACE2, by altering the proteolytic cleavages that occur at the S1-S2 boundary or by increasing stability of the spike protein. While the D614G mutation caused a highly significant increase in pseudotyped virus infectivity, the variants tested here, which include the mutation, provide either no increase in titer or a small increase (Δ69-70, S982A and Δ69-70/N501Y/P681H). However, the mutations could have greater effects on transmissibility as tissue culture does not model factors such as thermostability or stability in aerosols. In thermostability test, we found that the B.1.351 spike protein and some of the variant point mutations caused more than a 2-fold increase in infectivity at high temperature, an effect that may play a role in the increased transmissibility and prevalence of the variant viruses.

Our findings are consistent with recent reports from others. Wu *et al*. reported a decrease in titers of 2.7-fold for B.1.1.7 and 6.4-fold for the viruses with the B.1.351 spike protein by antibodies elicited by the Moderna 1273 mRNA vaccine^26^. Xie *et al*. found that BNT162b2 vaccine-elicited antibodies neutralized virus with the (E484K/N501Y) with a titer that was 0.8-fold compared to D614G^27^. Wang *et al*. found a more pronounced 11-33-fold decrease for convalescent serum neutralization of B.1.351 and 6.5-8.6-fold decrease for vaccinee sera^28^.

Spike protein-pseudotyped lentiviruses have been shown to provide an accurate measure of infectivity and antibody neutralization^29–34^. A comparison of neutralizing titers on a panel of 101 convalescent sera measured by lentiviral pseudotyped virus and live SARS-CoV-2 showed a high degree of concordance between the two assays^23^. This is the case even though the two assays are based on different viral particles and may not have similar numbers of spike protein trimers per virion. Whether this similarity is because pseudotyped viruses have a similar spike density to live virus or because neutralization sensitivity is independent of spike protein density is not clear.

The modest decrease in neutralizing titers against viruses with the B.1.1.7, COH.20G/677H and mink cluster 5 spike proteins by antibodies elicited by BNT162b2 vaccination suggests and the high titers induced by vaccination suggest that current vaccines are likely to be protective against currently circulating variant viruses. Viruses with the B.1.351 spike showed a 3-fold resistance to vaccine-elicited antibodies, but the given the high titers induced by vaccination, it seems likely that the vaccine will maintain a high degree of protection. However, one of the donor sera tested had relatively lower titer against B.1.351 suggesting that some individuals might not be highly protected. The decrease in titer was the result of the E484K mutation, suggesting that a modified vaccine with this mutation might be required to protect such individuals. Recent findings from in phase 3 vaccine trials by from Johnson and Johnson (https://www.jnj.com/johnson-johnson-announces-single-shot-janssen-covid-19-vaccine-candidate-met-primary-endpoints-in-interim-analysis-of-its-phase-3-ensemble-trial) and Novavax (https://ir.novavax.com/news-releases/news-release-details/novavax-covid-19-vaccine-demonstrates-893-efficacy-uk-phase-3) suggest a reduced level of protection from moderate to severe disease by the B.1.351 variant in South Africa.

As SARS-CoV-2 continues to rapidly evolve following its zoonotic transfer to humans, it is likely that novel variants will continue to emerge. These may have mutations in the spike protein that are selected both for escape from humoral responses and for increased transmissibility resulting from further increases in ACE2 affinity or increased stability. Our study, and others recently reported, demonstrate the importance of world-wide surveillance of circulating viruses by nucleotide sequencing and for monitoring novel spike proteins for neutralization by convalescent and vaccine sera. It will be important to define correlates of protection to understand the titer of neutralizing antibody required to provide protection against infection and disease pathogenesis and there is a need to produce modified vaccines based on variant mutations.

## Methods

### Plasmids

pLenti.GFP.NLuc is a dual GFP/nanoluciferase lentiviral vector based on pLenti.CMV.GFP.puro that contains a GFP/nanoluciferase cassette separated by a picornavirus P2A self-processing amino acid motif cloned into the BamH-I and Sal-I sites (Addgene plasmid #17448, provided by Eric Campeau and Paul Kaufman)^35^. pcCOV2.Δ19S is based on pCDNA6 in which the CMV promoter drives transcription of a chemically synthesized, codon-optimized SARS-CoV-2 spike gene based on the Wuhan-Hu-1/2019 amino acid sequence with a termination codon at position 1255 that results in the deletion of the carboxy-terminal 19 amino acids, as previously described^25^. Point mutations in the open reading frame were introduced by overlap extension. All plasmid sequences were confirmed by DNA nucleotide sequence analysis. HIV-1 Gag/Pol expression vector pMDL and HIV-1 Rev expression vector pRSV.Rev have been previously described^25^.

### Human sera

Convalescent sera and sera from individuals vaccinated at NYULH with BNT162b2 on day 0, day 7 and day 28 (7 days following the second injection) were collected from individuals through the NYU Vaccine Center with written consent under I.R.B. approval (IRB 20-00595 and IRB 18-02037) and were deidentified. Donor age and gender were not reported.

### Cells

293T cells were cultured in Dulbecco’s modified Eagle medium (DMEM) supplemented with 10% fetal bovine serum (FBS) and penicillin/streptomycin (P/S) at 37°C in 5% CO_2_. ACE2.293T cells are clonal cell-lines established by stable co-transfection with pLenti.ACE2-HA followed by selection in 1 μg/ml puromycin, as previously described ^23,25^.

### SARS-CoV-2 spike lentiviral pseudotypes

SARS-CoV-2 spike protein pseudotyped lentiviral stocks were produced by cotransfection of 293T cells with pMDL, pLenti.GFP-NLuc, pcCoV2.S-Δ19 (or variants thereof) and pRSV.Rev as previously described^25^. Virus stocks were normalized by real-time PCR reverse transcriptase activity^36^. Pseudotyped virus infections were done with 1 × 10^4^ cells/well in 96 well tissue culture dishes at an MOI=0.2 as previously described^25^. Luciferase activity was measured after 2 days using Nano-Glo luciferase substrate (Promega) and plates were read in an Envision 2103 microplate luminometer (PerkinElmer). To quantify antibody neutralization, sera were serially diluted 2-fold and then incubated for 30 minutes at room temperature with pseudotyped virus (corresponding to approximately 2.5 × 10^7^ cps luciferase) in a volume of 50 μl. The mixture was added to 1 × 10^4^ ACE2.293T cells (corresponding to an MOI of 0.2) in a volume of 50 μl in a 96 well culture dish. After 2 days, the medium was removed and Nano-Glo luciferase substrate (Nanolight) was added to wells. Luminescence was read in an Envision 2103 microplate luminometer (PerkinElmer).

### Immunoblot analysis

Cells were lysed in buffer containing 50 mM HEPES, 150 mM KCl, 2 mM EDTA, 0.5% NP-40, and protease inhibitor cocktail. Lysates (40 μg) were separated by SDS-PAGE and transferred to a polyvinylidene difluoride membrane. The membranes were probed with anti-spike mAb (1A9) (GeneTex), anti-p24 mAb (AG3.0), anti-His mAb (Invitrogen) and anti-GAPDH mAb (Life Technologies) followed by goat anti-mouse HRP-conjugated second antibody (Sigma). The membrane was treated with luminescent substrate (Millipore) and the band intensities were quantified on an iBright CL1000 Imager.

### Virus ACE2 binding assay

As previously described^25^, serially diluted recombinant soluble ACE2 protein containing carboxy-terminal His-tag were mixed with 20 μl Ni-NTA beads for 1 hour at 4°C. Unbound protein was removed by washing with PBS. The coated beads were then mixed with 40 μl pseudotyped lentiviral virions and incubated at 4°C for 1 hour. The beads were then washed with PBS, resuspended in reducing Laemmle loading buffer, heated to 90°C and separated by SDS-PAGE and analyzed on an immunoblot probed with anti-p24 antibody (AG3.0) followed by goat anti-mouse HRP-conjugated secondly antibody. The membrane was analyzed as described above.

### Soluble ACE2 Neutralization assay

As previously described^25^, serially diluted recombinant soluble ACE2 protein were mixed with pseudotyped virus for 1 hour at room temperature. The virus was added on the target cells and incubated for 2 days. After 2 days, the medium was removed and Nano-Glo luciferase substrate (Nanolight) was added to wells. The luminescence was read in an Envision 2103 microplate luminometer (PerkinElmer).

### Quantification and Statistical Analysis

All experiments were performed in technical duplicates or triplicates and data were analyzed using GraphPad Prism (Version 8). Statistical significance was determined by the two-tailed, unpaired t test. Significance was based on two-sided testing and attributed to p< 0.05. Confidence intervals are shown as the mean ± SD or SEM. (*P≤0.05, **P≤0.01, ***P≤0.001, ****P≤0.0001).

## Acknowledgements

We thank NYU Langone Vaccine Center staff Sara Hyman, Mahnoor Ali, Lisa Zhao, Heekoung Youn, Jimmy Wilson, Trishala Karmacharya, Joseph Allen, Sophie Gray-Galliard for their contributions. The work was funded by grants from the NIH to N.R.L. (DA046100, AI122390 and AI120898) and to M.J.M. (UM1AI148574), T.T. was supported by the Vilcek/Goldfarb Fellowship Endowment Fund.

## Author contributions

T.T. and N.R.L. conceived and designed the project. T.T. and B.M.D carried out the experiments and analyzed the data. M.S.G., R.S.H., A.C. and M.J.M. collected and provided the serum samples. T.T. and N.R.L wrote the manuscript. All authors provided critical comments on manuscript.

## Supplementary figures

**Supplementary Fig. 1.**
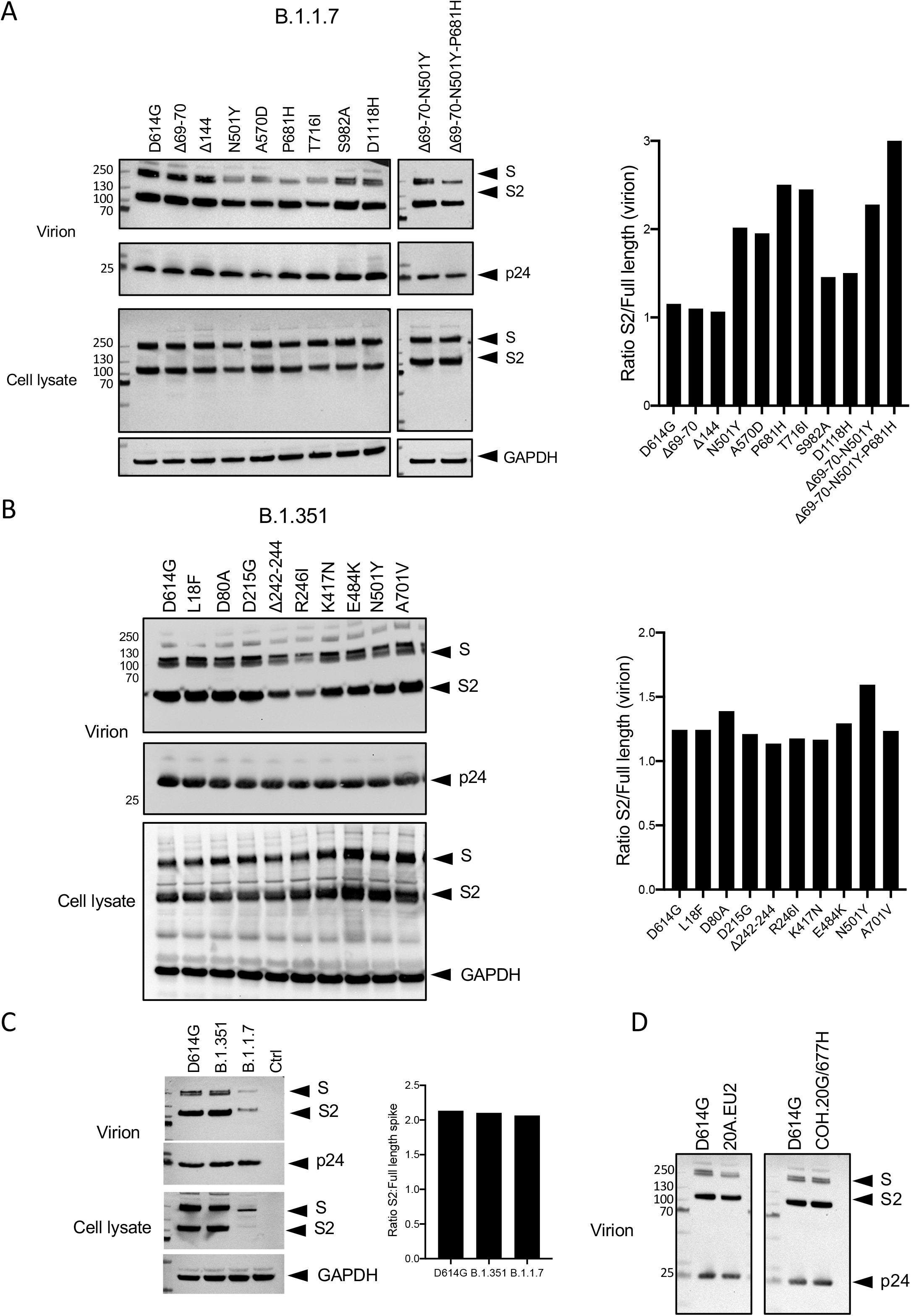
Immunoblot analysis of spike protein expression and incorporation into lentiviral particles. Pseudotyped viruses were generated by transfection of 293T cells. On day 2, cells and supernatant virions were collected, lysed and analyzed on an immunoblot probed with anti-His antibody to detect the full-length spike protein and S2 subunit. Cell lysates were probed with anti-GAPDH antibody to normalize for protein loading and virions were probed for p24 to normalize for virions. Arrows indicate the full-length spike (S), S2 subunit (S2), p24 and GAPDH. (A) Immunoblot analysis of spike proteins with individual B.1.1.7 mutations. Histogram on the right shows the ratio of full-length to S2 subunit determined by quantification of the bands. (B) Immunoblot analysis of spike proteins with individual B.1.351 mutations. Histogram on the right shows the ratio of S2 subunit to full-length spike protein determined by quantification of the bands. (C) Immunoblot analysis of spike proteins with D614G, B.1.1.7 and B.1.351 mutations. Histogram on the right shows the ratio of S2 subunit to full-length spike protein determined by quantification of the bands. (D) Immunoblot analysis of 20A.EU2 and COH20G/677H spike proteins.

**Supplementary Fig. 2.**
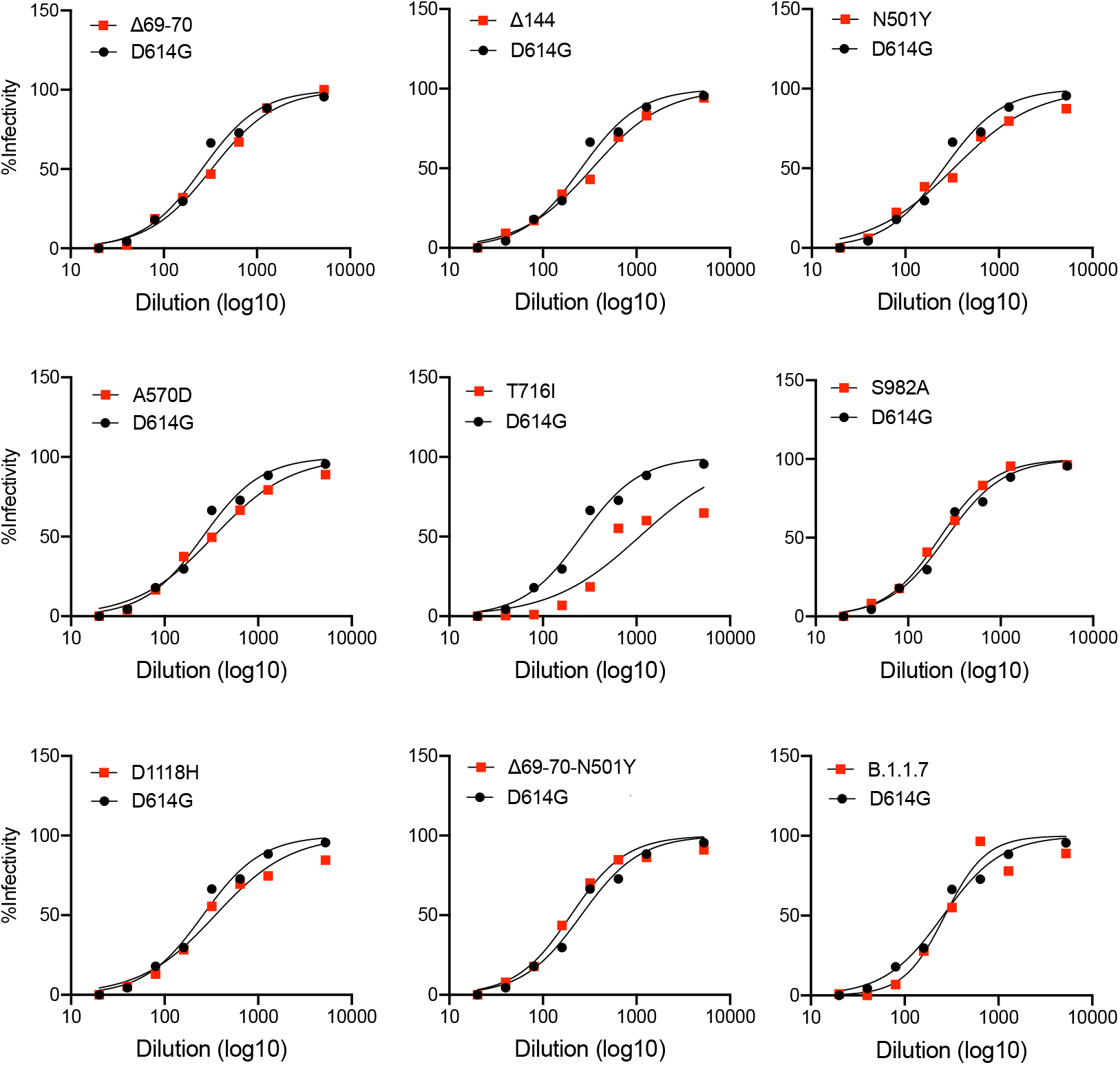
Viruses with B.1.1.7 are neutralized by convalescent serum. Convalescent serum neutralization titration curves of viruses with individual B.1.1.7 mutations.

**Supplementary Fig. 3.**
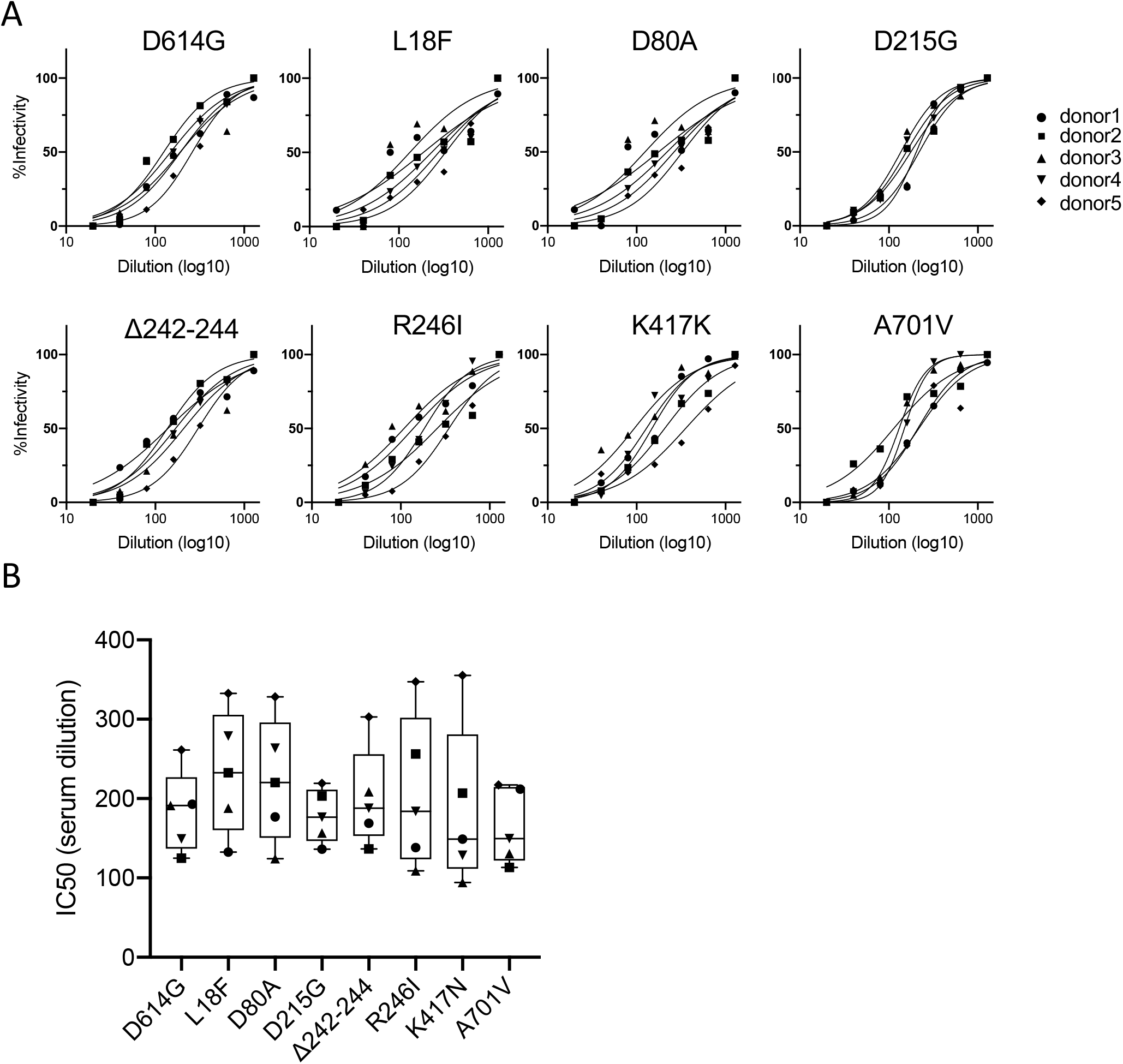
Convalescent serum neutralization titration curves of viruses with individual B.1.351 mutations. (A) Neutralization of viruses with individual B.1.351 mutations by sera of 5 convalescent individuals. The data represent percent infectivity not neutralized by convalescent sera. (B) Neutralization of viruses with the D614G and individual B.1.351 mutations by convalescent sera.

**Supplementary Fig. 4.**
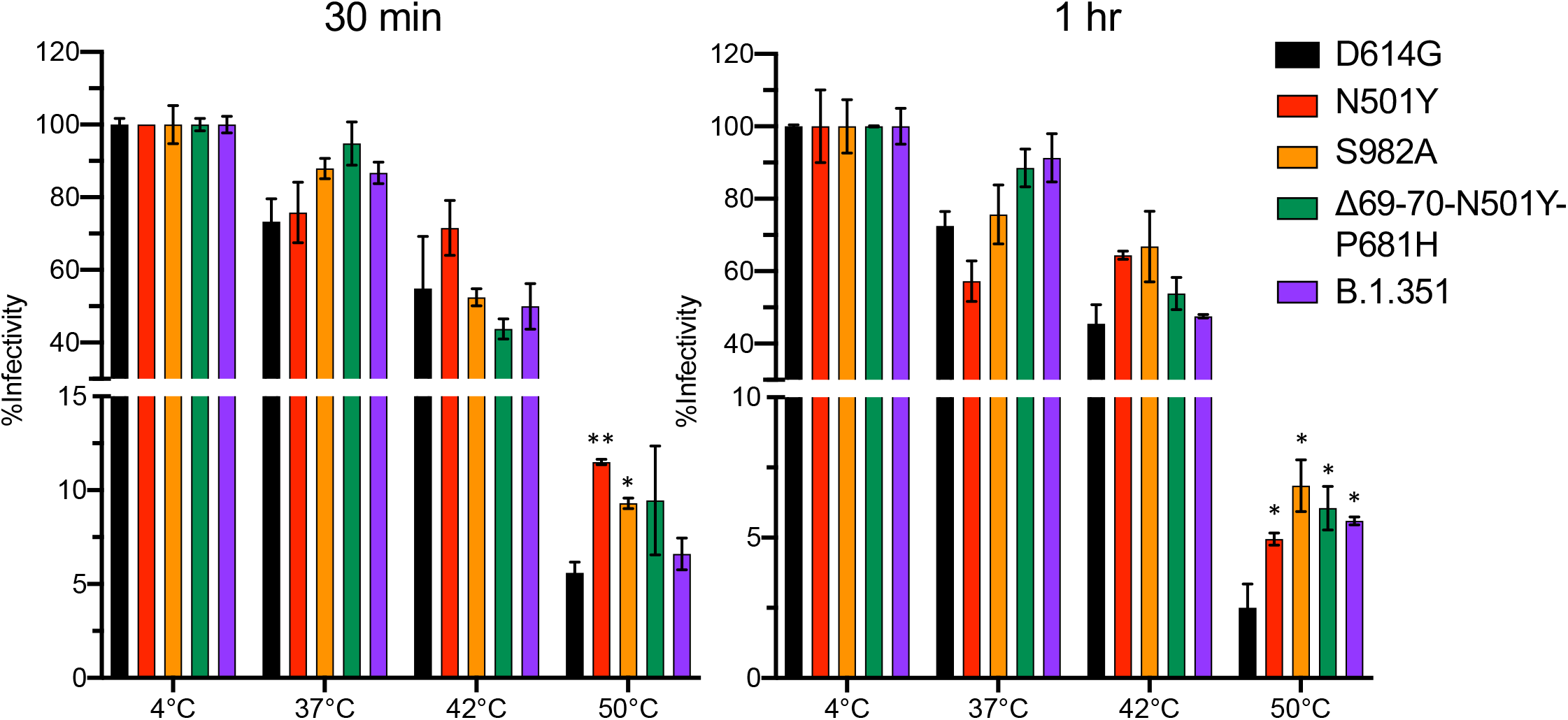
Thermostability of viruses with variant spike proteins. Viruses with the indicated spike proteins were incubated at 4°, 37°, 42° and 50°C for 30 min and 1 hour after which infectivity was determined in ACE2.293T cells.

